# It happened again – emergence of a Shamonda-like orthobunyavirus in Central Europe, 2026

**DOI:** 10.64898/2026.08.06.743243

**Authors:** Kerstin Wernike, Bernd Hoffmann, Ellen K. Link, Simon Rotheneder, Mirette Eshak, Florian Pfaff, Dirk Höper, Martin Beer

**Affiliations:** Friedrich-Loeffler-Institut, Südufer 10, 17493 Greifswald - Insel Riems, Germany; State Institute for Chemical and Veterinary Analysis Freiburg, Am Moosweiher 2, 79108 Freiburg, Germany; Animal Disease Fund Baden-Württemberg, Cattle Health Service Freiburg, Am Moosweiher 2, 79108 Freiburg, Germany

**Author notes:** **Address for correspondence:** Martin Beer, Friedrich-Loeffler-Institut, Federal Research Institute for Animal Health, Südufer 10, 17493 Greifswald - Insel Riems, Germany. These authors contributed equally to this article (Kerstin Wernike, Bernd Hoffmann).

**Keywords:** bunyavirus, cattle, ruminants, phylogeny, Simbu serogroup, Schmallenberg virus, Shamonda virus

## Abstract

Since mid-June 2026, an acute, non-fatal syndrome with high herd morbidity, reduced milk yield, diarrhea, fever and lethargy affected dairy cattle in southern Germany and Switzerland. Major viral pathogens were excluded. Pan-Simbuvirus PCR first detected an orthobunyavirus, subsequently confirmed by metagenomic sequencing, which enabled recovery of a complete “Shamonda-like” genome.

**One sentence summary line:** More than a decade after the emergence of Schmallenberg virus in 2011, a novel “Shamonda-like” orthobunyavirus of the Simbu serogroup has now emerged in Central Europe.

## Background

Since mid-June 2026, farmers and veterinarians in Southern Germany and Switzerland reported an acute disease in dairy cattle characterized by a marked decrease in milk production, diarrhea, fever, reduced activity and occasionally mild respiratory symptoms for a short period. By the end of July, well over 100 affected farms had been reported in the German federal state of Baden-Württemberg to the local cattle health service. At herd level, the morbidity rate was very high (up to 100%), but there have been no fatalities. Given the high temperatures at the time, heat stress was initially considered to be at least a contributing factor. On the other hand, an infectious disease was suspected, and a vector-borne virus was thought likely because of the time of year and the rate at which the disease was spreading. Classical endemic and emerging viruses such as bluetongue virus, epizootic hemorrhagic disease virus, Schmallenberg virus (SBV), pestiviruses, bovine herpesvirus type 1, bovine ephemeral fever virus, West Nil virus and Usutu virus could be excluded as the causative agent by (RT-)PCR.

In order to identify the cause of the outbreak, serum and EDTA blood samples from four herds in Baden-Württemberg (Figure 1) with acutely diseased cattle were sent by the local state laboratory to the Friedrich-Loeffler-Institut, Germany’s federal research institute for animal health, for further investigations using generic PCR systems and metagenomic analysis.

**Figure 1.**
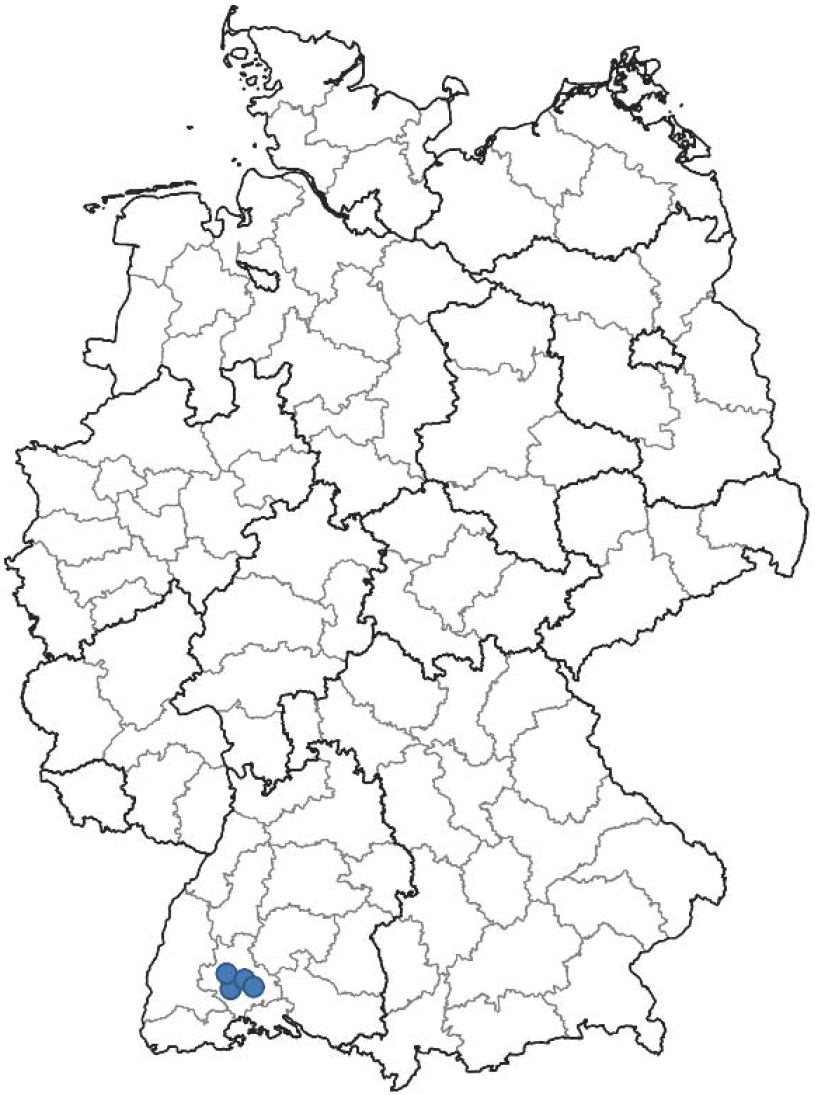
Location of cattle herds within Germany from which blood samples were subjected to metagenomic analysis and which tested positive for the Shamonda-like virus (blue dots). The boundaries of the German federal states (thick lines) and districts (thinner lines) are shown. The map was retrieved from the Federal Agency for Cartography and Geodesy (BKG; https://www.bkg.bund.de), data license Germany attribution version 2.0 (https://www.govdata.de/dl-de/by-2-0).

## This study

In total, 27 serum and 27 EDTA blood samples from four different cattle herds were submitted. Due to the small volume available per sample, up to six samples were pooled according to sample matrix and herd, resulting in 16 pools (two EDTA blood pools and two serum pools for each of the four herds). The samples were inoculated onto baby hamster kidney (BHK-21-BSR/5) and monkey kidney (Vero) cells (cell lines 0194 and 0228, Collection of Cell Lines in Veterinary Medicine, Friedrich-Loeffler-Institut, Germany) and infectious virus could be isolated from sample pools from three herds on Vero cells and from one of those pools additionally on BHK-21-BSR/5. Furthermore, nucleic acid was extracted from the sample pools with the King Fisher 96 Flex purification system and the NucleoMag Vet Kit (Machery Nagel) according to the manufacturer’s instructions. Subsequent testing by real-time RT-PCRs for ruminant orbiviruses, morbilliviruses, influenza A and D viruses, bovine ephemeral fever virus, and Hefer Valley virus gave consistently negative results. However, ten of the 16 pools tested positive by a generic real-time RT-PCR for viruses of the Simbu serogroup *(1)*, genus *Orthobunyavirus*, with quantification cycle (Cq) values ranging from 19.8 to 37.6. From every investigated cattle farm, at least one sample pool gave a positive result. Positive results were obtained from both sample matrices. In addition, all but 2 pools tested positive by an S-segment based real-time RT-PCR for Sathuperi virus (SATHV) *(2)* (Cq values 20.5 to 29.2). In contrast, all these samples tested negative by a specific real-time RT-PCR for SBV *(3)*, the only Simbu serogroup virus present in Central Europe so far *(4,5)*.

For metagenomic sequencing, RNA was extracted from TRIzol supernatants from serum and blood pools, respectively, using the RNAdvance Tissue Kit (Beckman Coulter) on a KingFisher Duo Prime platform (Thermo Fisher Scientific). For sequencing, the RNA was further processed as previously described *(6)*. Pooled libraries were loaded onto Ion 530 chips using an Ion Chef instrument and sequenced on an Ion S5 XL platform in 400-bp mode. The obtained datasets were assembled with Newbler (v3.0; Roche/454). The resulting complete genome sequence was submitted to the International Nucleotide Sequence Database Collaboration, http://www.insdc.org; project no. PRJEB123174. The obtained genome showed the highest identity to the Shamonda virus isolate Ib An 5550 (GenBank accession number HE795105.1), with 93% nucleotide identity for the L and M segment and 97% for the S segment. Hence, the virus was identified as a novel European Shamonda virus (SHAV) isolate within the species *Orthobunyavirus schmallenbergense*. Phylogenetic analyses were performed separately for each segment. Amino acid sequences of the proteins encoded by each segment were compared with 29 reference sequences obtained from the NCBI RefSeq database and aligned using MUSCLE v5.1. Maximum-likelihood trees were inferred with IQ-TREE v3.1.1 using the best-fit substitution model. Branch support was assessed with 10,000 ultrafast bootstrap and 10,000 SH-aLRT replicates. Sequences from the California and Bunyamwera serogroups were used as outgroups. All three genome segments were most closely related to Shamonda virus, providing no evidence for an inter-lineage reassortant origin (Figures 2, 3 and 4). In addition, the S and L segments were closely related to SBV (Figures 2 and 4), while the M segment differed markedly (Figure 3).

**Figure 2.**
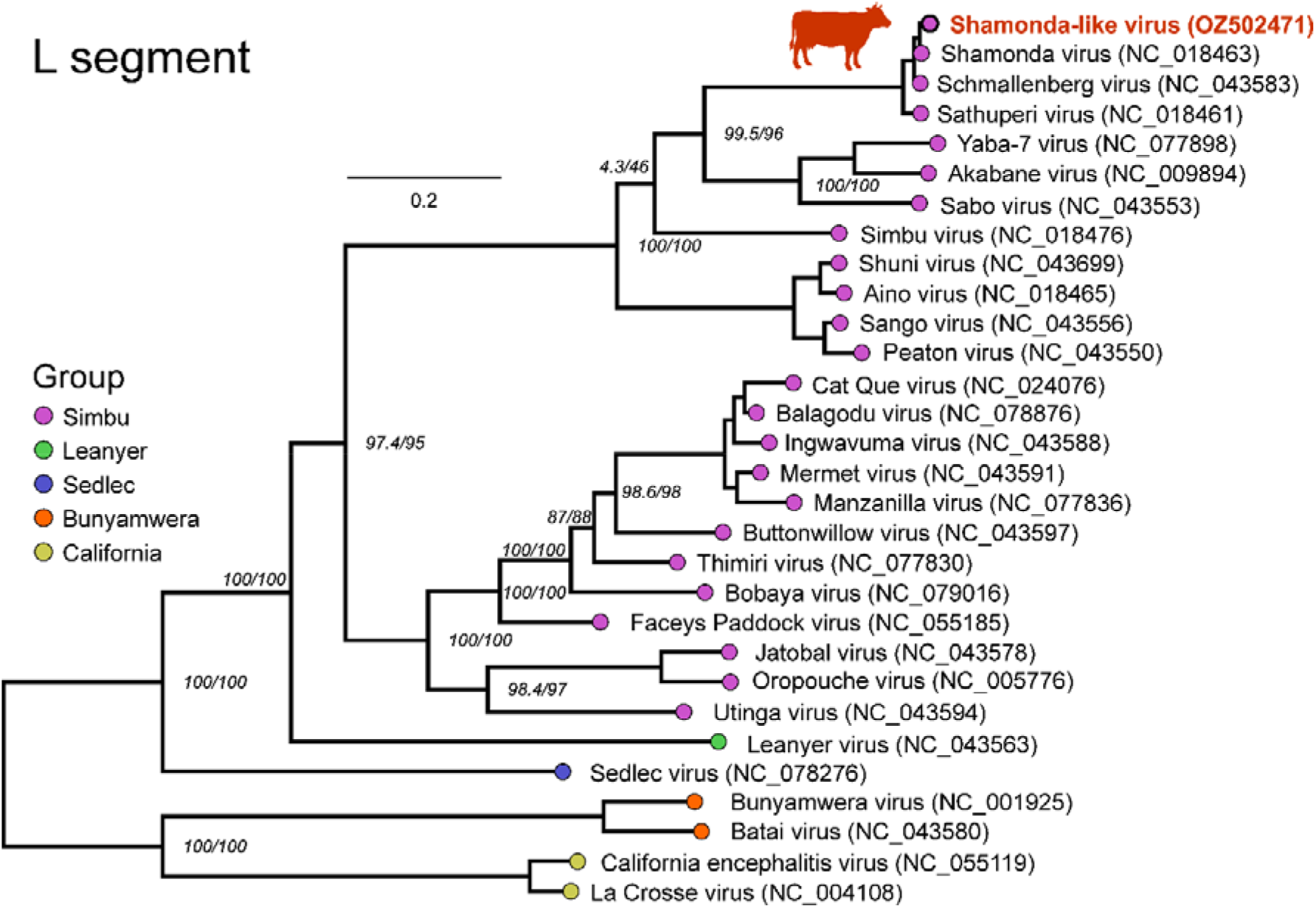
Phylogenetic analysis of the L segment of the novel Shamonda-like virus detected in Germany. The phylogenetic tree was inferred from the amino acid sequence of the RNA-dependent RNA polymerase encoded by the L segment of the novel Shamonda-like virus and 29 reference sequences. Sequences were aligned using MUSCLE v5.1, and maximum-likelihood phylogenetic analysis was performed with IQ-TREE v3.1.1 using the best-fit amino acid substitution model selected by the software. Branch support was assessed with 10,000 ultrafast bootstrap replicates and 10,000 SH-like approximate likelihood ratio test (SH-aLRT) replicates. Support values for major branches are shown in italics as ultrafast bootstrap/SH-aLRT percentages. Serogroups are indicated by color. Sequences belonging to the California and Bunyamwera serogroups were used as outgroups to root the tree.

**Figure 3.**
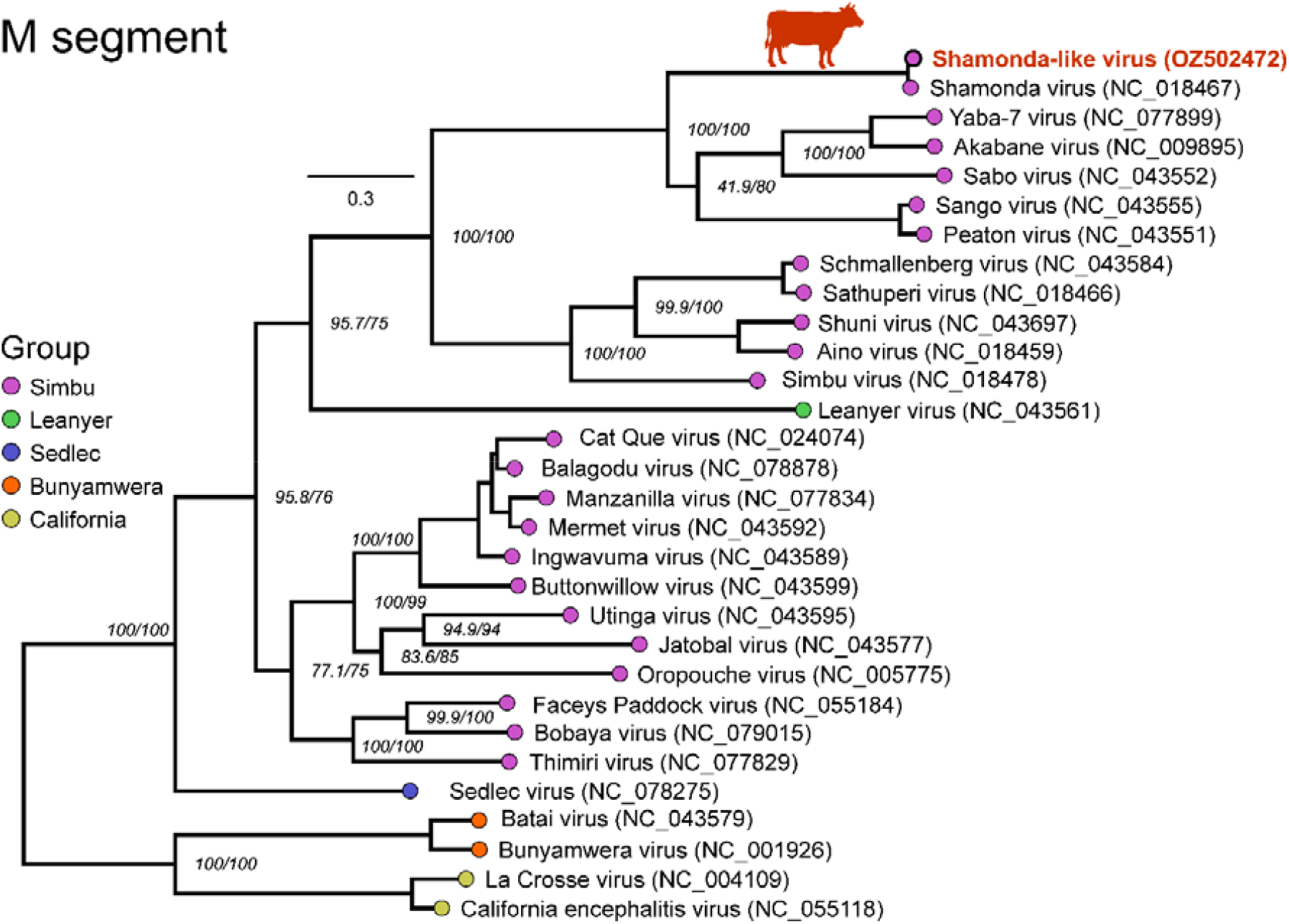
Phylogenetic analysis of the M segment of the novel Shamonda-like virus detected in Germany. The phylogenetic tree was inferred from the amino acid sequence of the glycoprotein precursor encoded by the M segment of the novel Shamonda-like virus and 29 reference sequences. Sequences were aligned using MUSCLE v5.1, and maximum-likelihood phylogenetic analysis was performed with IQ-TREE v3.1.1 using the best-fit amino acid substitution model selected by the software. Branch support was assessed with 10,000 ultrafast bootstrap replicates and 10,000 SH-like approximate likelihood ratio test (SH-aLRT) replicates. Support values for major branches are shown in italics as ultrafast bootstrap/SH-aLRT percentages. Serogroups are indicated by color. Sequences belonging to the California and Bunyamwera serogroups were used as outgroups to root the tree.

**Figure 4.**
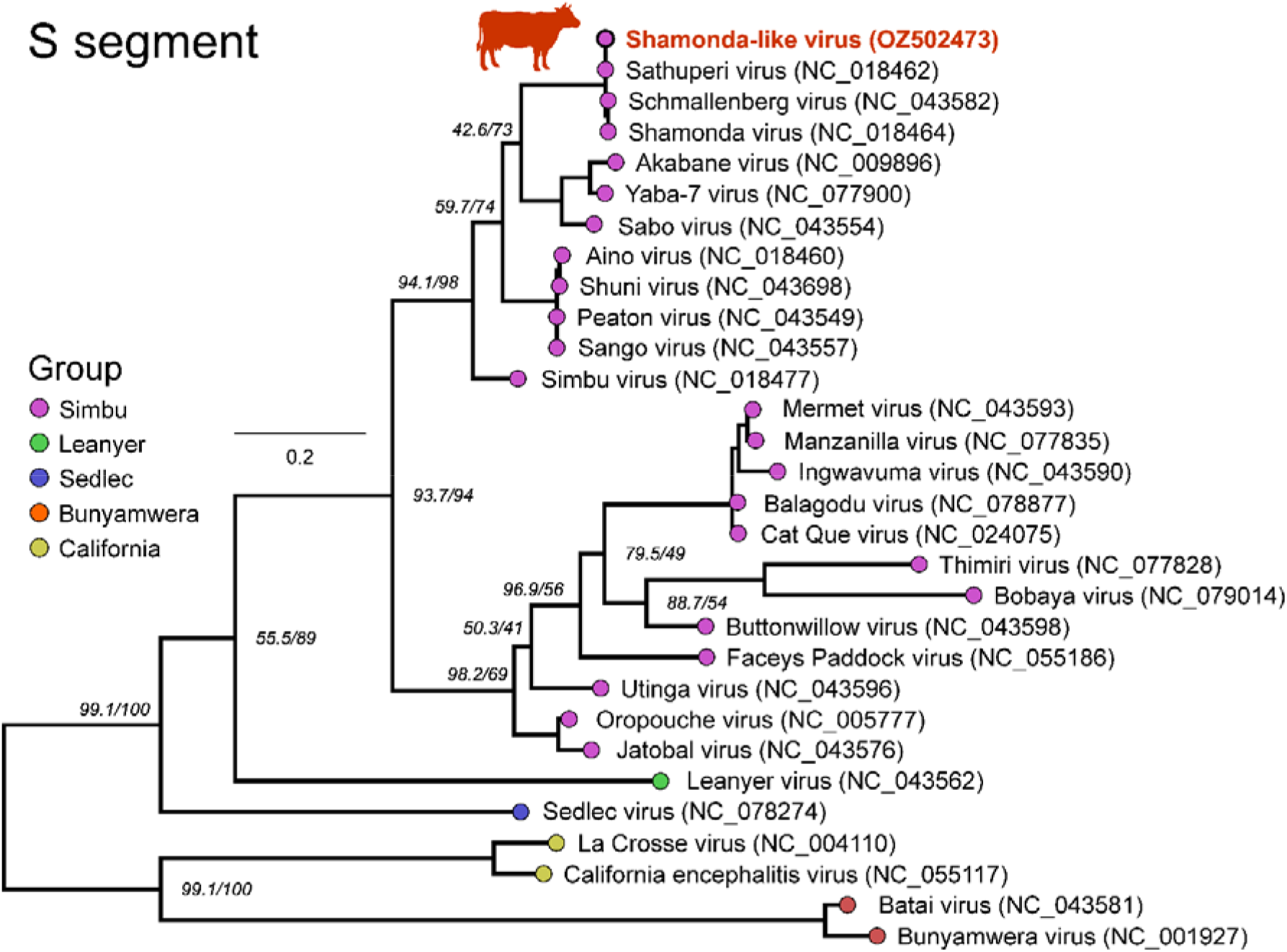
Phylogenetic analysis of the S segment of the novel Shamonda-like virus detected in Germany. The phylogenetic tree was inferred from the amino acid sequence of the nucleocapsid protein encoded by the S segment of the novel Shamonda-like virus and 29 reference sequences. Sequences were aligned using MUSCLE v5.1, and maximum-likelihood phylogenetic analysis was performed with IQ-TREE v3.1.1 using the best-fit amino acid substitution model selected by the software. Branch support was assessed with 10,000 ultrafast bootstrap replicates and 10,000 SH-like approximate likelihood ratio test (SH-aLRT) replicates. Support values for major branches are shown in italics as ultrafast bootstrap/SH-aLRT percentages. Serogroups are indicated by color. Sequences belonging to the California and Bunyamwera serogroups were used as outgroups to root the tree.

The emergence of a novel Simbu serogroup virus in the cattle population where SBV has been endemic for many years *(7)* may have important epidemiological and veterinary implications. Despite a high seroprevalence against SBV in the affected region, the large number of clinical cases associated with the newly introduced Shamonda-like virus strongly suggests that pre-existing immunity does not provide meaningful cross-protection. This observation is further supported by the sequence differences in the M-segment (Figure 4) which encodes the viral glycoproteins Gc and Gn, the major antigens of orthobunyaviruses *(8)*. Consequently, population immunity against SBV is unlikely to limit the spread of the Shamonda-like virus in the following summer and autumn months, assuming that the Shamonda-like virus is transmitted by biting insects, as are related simbuviruses *(4)*. Moreover, based on observations from other regions where two or more of the vector-transmitted simbuviruses are present, such as Australia or the Middle East, it is plausible that the two viruses will establish an alternating or cyclical pattern of circulation rather than one permanently displacing the other *(9,10)*. Such dynamics may be influenced by fluctuations in vector abundance, climatic conditions, and, above all, the level of herd immunity against one virus or the other.

Furthermore, reassortment of both viruses has to be expected in the future. Long-term surveillance will therefore be essential to determine whether similar epidemiological patterns will evolve in Europe and to identify factors driving the relative predominance of each virus over time.

At present, acute clinical disease has only been reported in cattle in Germany, France and Switzerland. However, the host range of the newly emerged European Shamonda-like virus remains incompletely understood. Though for SHAV infections have mainly been documented in cattle so far *(11,12)*, infection of other domestic and wild ruminant species cannot be excluded, given the broader range of ruminant hosts observed for several other members of the Simbu serogroup, including SBV *(7)*. Therefore, monitoring sheep, goats, and free-ranging ruminants during the coming months will be crucial to assess the host range. In this context, it is particularly important to monitor animals that were pregnant at the time of infection, as a particularly worrying feature of multiple simbuviruses including SHAV and SBV is that they can cause severe fetal malformations, when naïve dams are infected during early gestation *(3,4,10,12)*. Such investigations will not only improve understanding of the pathogenesis and ecology of the virus but will also inform risk assessments. Continued molecular and serological surveillance across susceptible species, combined with vector monitoring and identifying the local vector species responsible for virus transmission, will be critical for characterizing the long-term epidemiology of the emerging Shamonda-like virus and for guiding appropriate disease control measures.

## Acknowledgments

We thank Tobias Töws, Patrick Zitzow, Karin Pinger and Bianka Hillmann for excellent technical assistance.

## Ethical statement

The samples were taken by veterinarians in the context of health monitoring and as part of their clinical practice, no permissions were needed to collect these specimens.

## About the Authors

PD Dr. Kerstin Wernike and Dr. Bernd Hoffmann are veterinarians and senior scientists at the Friedrich-Loeffler-Institut, Greifswald-Insel Riems, Germany. Their research interests focus on emerging viruses, host-virus-interaction, diagnostics and immunoprophylaxis.

## References

1. Golender N, Bumbarov VY, Erster O, Beer M, Khinich Y, Wernike K. Development and validation of a universal S-segment-based real-time RT-PCR assay for the detection of Simbu serogroup viruses. J Virol Methods. 2018 Nov;261:80–5.

2. Tauscher K, Wernike K, Fischer M, Wegelt A, Hoffmann B, Teifke JP, Beer M. Characterization of Simbu serogroup virus infections in type I interferon receptor knock-out mice. Arch Virol. 2017 Oct;162(10):3119–29.

3. Bilk S, Schulze C, Fischer M, Beer M, Hlinak A, Hoffmann B. Organ distribution of Schmallenberg virus RNA in malformed newborns. Vet Microbiol. 2012 Sep 14;159(1-2):236–8.

4. Sick F, Beer M, Kampen H, Wernike K. Culicoides biting midges-underestimated vectors for arboviruses of public health and veterinary importance. Viruses. 2019 Apr 24;11(4).

5. Hoffmann B, Scheuch M, Höper D, Jungblut R, Holsteg M, Schirrmeier H, et al. Novel orthobunyavirus in cattle, Europe, 2011. Emerg Infect Dis. 2012 Mar;18(3):469–72.

6. Eschbaumer M, Staubach C, Pfaff F, Gethmann J, Schulz K, Rogoll L, et al. Buffaloed in Brandenburg: Germany’s first Brush with Foot-and-Mouth Disease after four Decades of Freedom. bioRxiv. 2026.

7. Wernike K, Beer M. More than a decade of research on Schmallenberg virus - Knowns and unknowns. Adv Virus Res. 2024;120:77–98.

8. Hellert J, Aebischer A, Wernike K, Haouz A, Brocchi E, Reiche S, et al. Orthobunyavirus spike architecture and recognition by neutralizing antibodies. Nat Commun. 2019 Feb 20;10(1):879.

9. O’Connor TW, Hick PM, Toribio J, Peel AJ, Kirkland PD, Finlaison DS. The Dynamic Transmission of Simbu Group Viruses in New South Wales, Australia. Transbound Emerg Dis. 2026;2026(1):e9214955.

10. Golender N, Bumbarov V, Kovtunenko A, David D, Guini-Rubinstein M, Sol A, et al. Identification and genetic characterization of viral pathogens in ruminant gestation abnormalities, Israel, 2015-2019. Viruses. 2021 Oct 22;13(11):2136.

11. Causey OR, Kemp GE, Causey CE, Lee VH. Isolations of Simbu-group viruses in Ibadan, Nigeria 1964-69, including the new types Sango, Shamonda, Sabo and Shuni. Ann Trop Med Parasitol. 1972 Sep;66(3):357–62.

12. Hirashima Y, Kitahara S, Kato T, Shirafuji H, Tanaka S, Yanase T. Congenital malformations of calves infected with Shamonda virus, Southern Japan. Emerg Infect Dis. 2017 Jun;23(6):993–6.

